# In vivo analysis of the evolutionary conserved BTD-box domain of Sp1 and Btd during Drosophila development

**DOI:** 10.1101/2020.03.25.007625

**Authors:** David Blom-Dahl, Sergio Córdoba, Hugo Gabilondo, Pablo Carr-Baena, Fernando J. Díaz-Benjumea, Carlos Estella

**Author notes:** These authors have contributed equally to this work (David Blom-Dahl, Sergio Córdoba).

## Abstract

The Sp family of transcription factors plays important functions during development and disease. An evolutionary conserved role for some Sp family members is the control of limb development. The family is characterized by the presence of three C2H2-type zinc fingers and an adjacent 10 aa region with an unknown function called the Buttonhead (BTD) box. The presence of this BTD-box in all Sp family members identified from arthropods to vertebrates, suggests that it plays an important role during development. However, despite its conservation, the *in vivo* function of the BTD-box has never been studied. In this work, we have generated specific BTD-box deletion alleles for the *Drosophila* Sp family members *Sp1* and *buttonhead* (*btd*) using gene editing tools and analyzed its role during development. Unexpectedly, *btd* and *Sp1* mutant alleles that lack the BTD-box are viable and have almost normal appendages. However, in a sensitized background the requirement of this domain to fully regulate some of Sp1 and Btd target genes is revealed. Furthermore, we have also identified a novel Sp1 role promoting leg *vs* antenna identity through the repression of *spineless* (*ss*) expression in the leg, a function that also depends on the Sp1 BTD-box.

## Introduction

The evolutionary conserved family of Sp transcription factors has been implicated in multiple developmental processes and diseases from *C. elegans* to humans [1, 2]. Sp proteins bind to GC rich motifs in promoters and enhancers regulating the transcription of their target genes. The common structural features of all Sp members are the presence at the C-terminal region of three C2H2-type zinc fingers and the Buttonhead (BTD) box [3, 4]. First identified in the *Drosophila* gene *buttonhead* (*btd*) [5], the BTD-box is a 10 aa stretch (R-X0–4-C-X-[C/D/N]-P-[N/Y]-C) adjacent to the DNA binding domain with function is unknown. The existence of the BTD-box in all Sp family members identified from arthropods to vertebrates suggested an important role during development compared to the more variable presence of other structural domains [3]. Importantly, deletion of a domain that contains the BTD box in the human Sp1 suggested a transactivation role in *in vitro* experiments [6–8]. Importantly, despite its conservation, the *in vivo* requirements of the BTD-box have never been studied.

An important and evolutionary conserved function of the members of the Sp family is their role during appendage development across many animal phyla. Although non-homologous structures, vertebrate and arthropod appendages share a similar and ancient developmental program [9]. Genes like *Distalless* (*Dll/Dlx*), *homothorax* (*hth/meis*) and members of the Sp family are expressed and required for appendage formation from mice to flies [10–16]. The *Drosophila* genome encodes for three members of the Sp family: *Spps* (*Sp1-like factor for pairing sensitive-silencing*), *btd* and *Sp1* [3]. Of these, only *btd* and *Sp1* have been reported to be required for leg development [13, 17, 18]. These transcription factors also play important roles in *Drosophila* such as embryo head segmentation (only for *btd*) and type II neuroblast (NB) specification and maintenance (*btd* and *Sp1*) [5, 19, 20].

In *Drosophila*, appendage formation is initiated during embryogenesis by the specification of a group of cells that form the thoracic imaginal primordia (reviewed in [21]). These cells activate the expression of *Dll* and are the progenitors of the ventral (leg) and dorsal appendages (wing and haltere) [22–24]. The leg identity is established in the embryo by the activity of Btd and Sp1 that act redundantly to induce *Dll* expression trough dedicated *cis*-regulatory modules (CRMs) [13, 17]. During larval stages, the leg imaginal primordium grows and is progressively patterned by the combined action of signaling pathways and transcription factors [21, 25]. The leg proximo-distal (PD) axis is initiated by the joint action of the Wingless (Wg) and Decapentaplegic (Dpp) morphogenes that regulate the expression of *Dll*, *dachsound (dac)* and *hth* in distal, medial and proximal domains, respectively [25–28]. In addition, another signaling pathway, the Epidermal Growth Factor Receptor (EGFR) patterns the distal-most region of the disc through the activation of a specific set of transcription factors that ensure the segmental subdivision of the tarsal domain in five tarsi [29–32]. The combinatorial code of transcription factors expressed along the PD axis of the leg is ultimately responsible for the localization of the Notch ligands Delta (Dl) and Serrate (Ser) [33]. Notch pathway activation in concentric rings at the distal end of each segment directs the formation of the joints and controls the growth of the leg [34–36] and reviewed in [37].

During leg imaginal development, *btd* and *Sp1* have both similar and diverging functions that are reflected by their dynamic expression patterns [18]. During embryogenesis, both genes are expressed in the thoracic ventral limb (leg) primordia and later, throughout larval development their expression diverge, being Sp1 restricted to the distal leg while *btd* extending more proximally [3, 13, 17, 18]. Initially, both genes act redundantly to activate *Dll* together with Wg and Dpp and only the elimination of both *btd* and *Sp1* abolish *Dll* expression [17, 18]. Once *Dll* is tuned on, its expression is maintained, in part, through an autoregulatory mechanism and no longer depends on *Sp1* or *btd* [27, 38]. At this time, the function of *btd* and *Sp1* is co-opted to regulate the growth and morphogenesis of the leg. The leg phenotypes generated by *btd* and *Sp1* specific mutations reveal the different contributions of each gene to appendage development. Elimination of *btd* from the entire leg mostly affected the growth of the proximo-medial segments while *Sp1* mutants presented dwarfed legs with segment fusions [17, 18]. Molecular analysis of Sp1 function demonstrated that Sp1 activates the expression of the Notch ligand *Ser* in the tarsal segments.

In addition, through a transcriptome analysis of *Sp1* mutant legs, several Sp1 target genes were identified. Interestingly, some of these genes are the antenna-specific gene *distal antenna-related* (*danr*) and *spineless* (*ss*) that are upregulated in *Sp1* mutant legs suggesting a role for Sp1 repressing antennal fates [18]. Leg and antenna are homologous appendages that share a common patterning logic but differ in the expression of the homeotic gene *Antennapedia* (*Antp*) that specifies leg *versus* antenna identity [39]. In the leg, the expression of *Dll* and *hth* is mostly exclusive, in part though the repression of *hth* by Antp. However, in the antenna the broad co-expression of both genes is required for antennal specification [40–42]. In the antenna, the combined action of Dll and Hth activate the expression of *ss* and the antenna-specific gene *spalt* (*sal*) [40, 43–45]. In particular, *ss* encodes for a bHLH-PAS transcription factor related to the mammalian dioxin receptor [46]. *ss* is expressed and required from the 3^rd^ antennal segment to the distalmost segments, while in the leg is only transiently expressed and it is necessary to specify the tarsal primordia [46]. In some *ss* mutant conditions, there is a transformation of the distal antenna into tarsal segments and its ectopic expression in the leg induced the formation of antennal fates [46]. Ss activate the expression of the sister genes *distal antenna* (*dan*) and *danr* that contribute to the specification of distal antenna structures [47, 48]. An important and unresolved question is how Sp1/Btd repress the antennal developmental program in the leg.

In this study, we first investigated the role of the BTD-box by generating *Sp1* and *btd* mutant alleles that lack this specific domain. In addition, we complemented these experiments with phenotypic analysis of gain of function of *Sp1* and *btd* versions without the BTD-box. Second, we described the molecular mechanism employed by Sp1 to repress antennal fates in the leg. Unexpectedly *Sp1* and *btd* mutant flies lacking the BTD-box domain were viable and produced normally patterned appendages of nearly the right size. However, in a sensitized background, the lack of the BTD-box in *Sp1* and *btd* reveals an important role modulating their transcriptional output. We also analysed the role of the BTD-box in other tissues in which it has been reported that Btd and Sp1 play a function, like the specification of the head segments in the embryo and the maintenance of type II NB identity in the larval brain. In addition, we have also identified *ss* as a direct target of Sp1 in the leg. Taken together, our results provide important information about the molecular mode of action of Sp1 and Btd during development.

## Results

### Generation of *Sp1* and *btd* BTD-box deletion alleles

To analyze the possible *in vivo* role of the BTD-box conserved domain, we have taken advantage of the *Sp1*^*HR*^ null allele that we have previously generated, which presents an *attP* integration site in place of the third *Sp1* exon [18] (Fig. 1A). To this end, we generated and inserted back a version of the *Sp1* exon that lacks the BTD-box (*Sp1* ^*ΔBTD-box*^) (Fig. 1A). As a control, we restored the wild type third exon (*Sp1*^*Wt*^) and confirmed its ability to completely rescue the *Sp1*^*HR*^ leg phenotypes (see below). In addition, we have generated, through CRISPR/Cas9 technology, a 4 aa and a 6 aa deletion of the BTD-box domain of *btd* in a wild type chromosome (named *btd*^*ΔBTD-box*^) and over the *Sp1*^*ΔBTD-box*^ allele (named *Sp1*^*ΔBTD-box*^, *btd*^*ΔBTD-box*^), respectively (Figure 1A).

**Figure 1:**
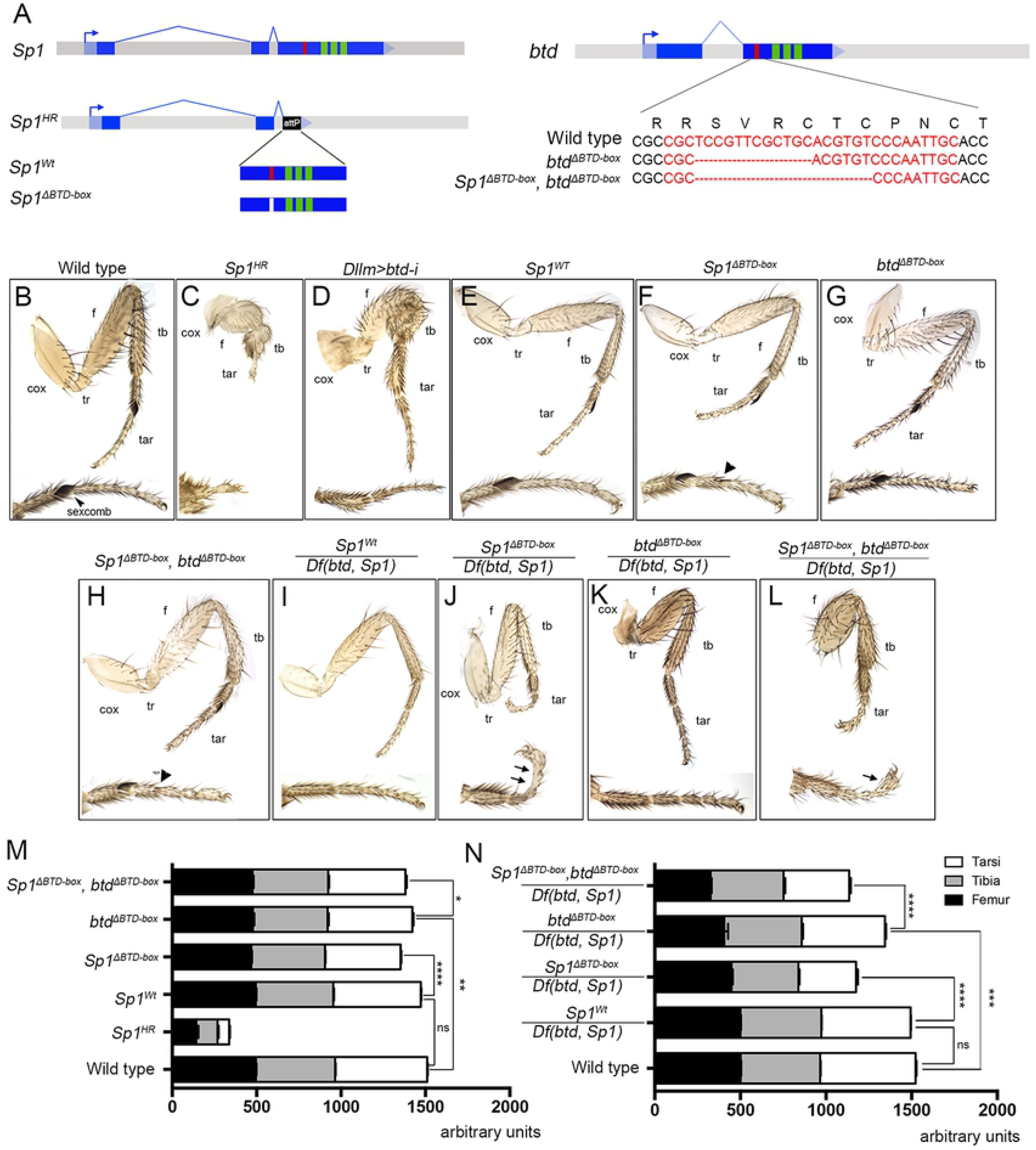
Generation of *Sp1* and *btd* BTD-box deletion alleles and adult phenotypes. (A) Schematic representation of the *Sp1*^*Wt*^, *Sp1*^*ΔBTD-box*^, *btd*^*ΔBTD-box*^ and the double *Sp1*^*ΔBTD-box*^, *btd*^*ΔBTD-box*^ alleles. The attP integration site (black box) that substitutes the third exon of *Sp1* in the *Sp1*^*HR*^ null allele was used to integrate a wild type copy (*Sp1*^*Wt*^) or a BTD-box deletion version (*Sp1*^*ΔBTD-box*^) of the exon. Two *btd*^*ΔBTD-box*^ alleles were generated by CRISPR/Cas9 in a wild type chromosome (4 aa deletion) and in the *Sp1*^*ΔBTD-box*^ background (6 aa deletion). Within the third exon, the three zinc fingers (green) and the BTD- box (red) are highlighted. (B-L) Adult leg phenotypes for the different genetic combinations described in the main text and above each image. Below each leg, a detailed view of the tarsal region is shown. cox, coxa; f, femur; tar, tarsus; tb, tibia; tr, trochanter. Arrowhead indicate the presence of ectopic sexcomb like bristles in F and H and arrows point to joint defects in J and L. (M and N) Leg size quantification for the different genetic combinations shown in B to L. In M only first legs from males were quantified while in N only first legs from females were measured. n>20 except from *btd*^*ΔBTD-box*^/*Df(btd,Sp1)* (n=14) and for *Sp1*^*ΔBTD-box*^, *btd*^*ΔBTD-box*^/*Df(btd,Sp1)* (n=10). **** P ≤ 0.0001, *** P ≤ 0.001, ** P ≤ 0.01, * P ≤ 0.05 with Student’s t test indicating a significant difference in each comparison. ns, non-significant. Error bars represent standard error of the mean (SEM).

### *Sp1* and *btd* BTD-box deletion alleles present defects in leg development in a sensitized background

*Sp1*^*HR*^ mutant flies die as pharate, and have strong defects in leg growth and joint formation due to the requirement of Sp1 to regulate Notch activity (Fig. 1C) [18]. *btd* null mutants die during embryogenesis with defects on head segmentation [5, 49]. However, leg-specific knockdown of Btd levels by the expression of a *btd-RNAi* (*Dllm>btd-i*, see Material and Methods) cause strong growth defects, more evident in the femur and tibia that sometimes appear fused (Fig. 1D) [17, 18]. *Sp1*^*HR*^ defects on leg growth and joint formation were completely rescued in the restored *Sp1*^*Wt*^ allele (compare Fig. 1E with 1C and 1M for quantification). Unexpectedly due to Btd-Box conservation in all Sp members, deletion of the entire domain in *Sp1*, or a partial deletion in *btd*, doesn’t cause major defects on fly viability or patterning (Fig. 1F and 1G). Nevertheless, a detailed characterization of *Sp1*^*ΔBTD-box*^ adult legs shows that, compared to the *Sp1*^*Wt*^ control, deletion of the BTD-box cause subtle, although consistent, leg size defects that are more evident in the tarsal region (Fig. 1F and 1M). In addition, we observed a decrease in the number of bristles that form the sexcombs (≈8.41 in the mutant *vs* ≈11.65 in the *Sp1*^*Wt*^ control, n>20) and the appearance of extra sexcomb-like bristles in more distal segments (Figure 1F). Similarly, *btd*^*ΔBTD-box*^ animals are viable and normally patterned with a very modest leg size reduction (compare Fig. 1G with 1B and 1M). As *btd* and *Sp1* have partially redundant functions during leg development [13, 17], we tested the combined deletion of the BTD-box in a *btd*^*ΔBTD-box*^, *Sp1*^*ΔBTD-box*^ double mutant chromosome. The viability of these mutant flies is compromised, however the few adult homozygous mutant flies recovered have similar leg-patterning defects as *Sp1*^*ΔBTD-box*^ mutants (Figure 1H and 1M).

Both in *Drosophila* and in mouse, a progressive reduction in the dose of Sp members increases the severity of limb phenotypes [17, 18, 50]. Therefore, we tested the function of the BTD-box domain of *Sp1* and *btd* in a sensitized background by decreasing the dose of these genes using a deficiency for both of them (*Df(btd, Sp1)*) [17]. As these genes are located in the X chromosome, we were only able to analyze females of the desired genotypes. Animals that have only one functional allele for *btd* and the *Sp1*^*ΔBTD-box*^ mutant allele (*Sp1*^*ΔBTD-box*^/*Df(btd, Sp1*)) present strong tarsal joint defects and an overall reduction in leg size that is more apparent in the distal region, as compared to the experimental control (*Sp1*^*Wt*^/*Df(btd, Sp1*)) that has a wild type phenotype (compare Fig. 1J with 1I and 1N). Animals that have the *btd*^*ΔBTD-box*^ allele over the *Df(btd, Sp1)* die during larval and pupal development, however very few escapers (less than 1%) can be recovered that die as pharate. These escapers also present an obvious leg size reduction, which is more apparent in the femur that in a few cases appears fused to the tibia (Fig. 1K). Interestingly, the defects observed for the *Sp1* and *btd* BTD-box mutant alelles over the *Df(btd, Sp1)* reproduce, to some extent, the phenotypes described for the single mutants of these genes [17, 18](Figure 1C and 1D). We were only able to obtain very rare escapers of the double *btd*^*ΔBTD-box*^, *Sp1*^*ΔBTD-box*^ mutant chromosome over the *Df(btd, Sp1)* that show strong defects on leg growth and joint formation (Fig. 1L and 1N).

All together, these results suggest a role for the BTD-Box during leg development that is better evidenced when the dose of the *Sp1* and *btd* genes is compromised.

### The BTD-box of *Sp1* is required for *Ser* expression in the distal leg

In the leg, the Notch pathway directs the formation of the joints and controls the growth of the appendage [35, 36]. Notch activation in concentric rings depends on the precise localization of the Notch ligands, Ser and Dl, at the distal end of each segment, which is regulated by different PD transcription factors [33]. We have previously shown that Sp1, in combination with other PD transcription factors, regulate *Ser* expression in the distal leg [18]. Therefore, we analyzed *Ser* expression and Notch activation in the different mutant combinations that lack the BTD-box.

*Sp1*^*HR*^ mutant prepupal leg imaginal discs are characterized by a strong decrease in Ser levels in the tibia and the tarsi accompanied by a reduction of the rings of the Notch target gene *big brain* (*bib*) (compare Figure 2A with 2B). Accordingly to the adult phenotypes, no apparent defects were observed in the expression of *Ser* and *bib* in *Sp1*^*ΔBTD-box*^ or *btd*^*ΔBTD-box*^ mutant animals (Figure 2C, D and E). Importantly, when using a sensitized background, incomplete rings of Ser and *bib* were observed in the tarsal region of *Sp1*^*ΔBTD-box*^/*Df(btd, Sp1*) prepupal leg discs when compared to the experimental control (compare Figure 2G and F). No defects were appreciable in *Ser* and *bib* levels in discs of the *btd*^*ΔBTD-box*^/*Df(btd, Sp1*) genotype (Figure 2H), which is consistent with the lack of tarsal joint defects in this genotype (see Figure 1K). Unfortunately, we were not able to dissect *Sp1*^*ΔBTD-box*^, *btd*^*ΔBTD-box*^/*Df(btd, Sp1*) animals due to the low frequency of escapers. In summary, these results suggest a role for the BTD-Box domain in the regulation of Notch activity during leg development in conditions where the dose of *Sp1* and *btd* is reduced.

**Figure 2:**
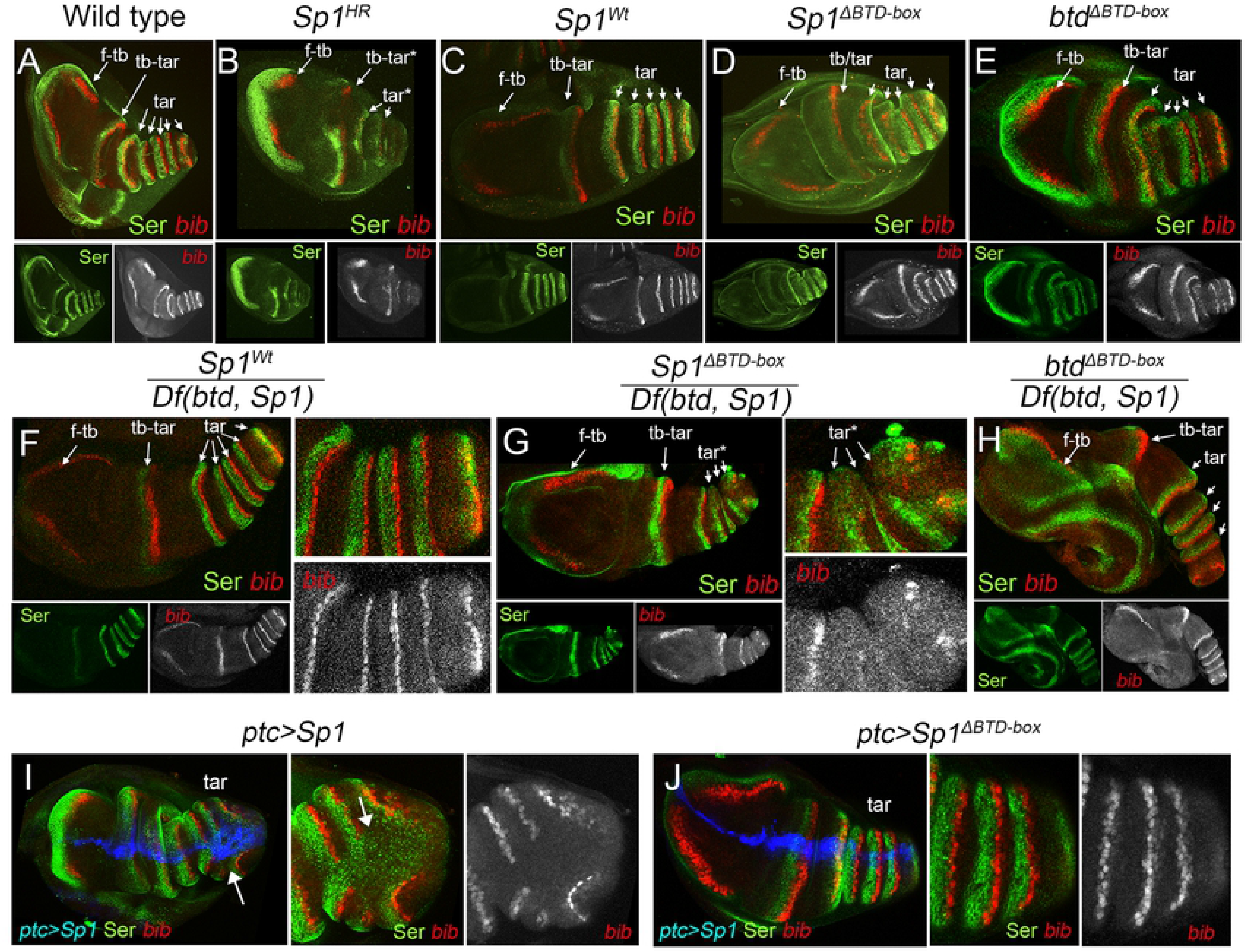
The BTD-box of *Sp1* is required for correct *Ser* expression in the distal leg. (A-H) Ser staining (green) and *bib-lacZ* (red) expression in prepupal legs from the different genotypes indicated above each panel. Separate channels for Ser and *bib* stainings are shown below. f-tb, femur-tibia joint; tb-tar, tibia-tarsal joint; tar, tarsal joints. Note the absence or incomplete formation of *Ser* and *bib* rings, marked by asterisks, in the *Sp1*^*HR*^ and *Sp1*^*ΔBTD-box*^/*Df(btd,Sp1)* mutant legs. Compare B with A and G with F panels. Detailed view of the tarsal region is shown for F and G. (I and J) Prepupal leg discs stained for Ser (green) and *bib-lacZ* (red) where UAS-*Sp1* (I) or UAS-*Sp1*^*ΔBTD-box*^ (J) were ectopically expressed starting from third instar larval stage using the *ptc-Gal4, UAS-GFP*; *tubGal80*^*ts*^ driver (blue). White arrow indicates misexpression of *Ser* that was restricted to the tarsal region. Detailed view of the tarsal region is shown for I and J. Note that the ectopic expression of *Sp1*, but not of *Sp1*^*ΔBTD-box*^, activates *Ser* expression and represses *bib-lacZ*.

### Defect on type II neuroblast specification in *Sp1* and *btd* BTD-box deletion alleles

Interestingly, individuals carrying either the double *Sp1*^*ΔBTD-box*^, *btd*^*ΔBTD-box*^ mutant allele or the different combinations of the BTD-box mutants over the *Df(btd, Sp1*) rarely pass pupal stages, suggesting a role of the BTD box for animal viability. Therefore, we decided to study the requirement of this domain for other known functions executed by Sp1 and Btd. Besides their role during appendage development, *btd* controls the formation several head segments (intercalary, antennal and mandibular) [49], while both *btd* and *Sp1* are required for type II NB identity specification [20]. First, we investigated the expression of *Dll* as a marker of the antennal segment and ventral thoracic primordia in *btd*^*ΔBTD-box*^/*Df(btd, Sp1*) mutant embryos. In contrast to the *btd*^*XG*^ null mutant that lacks the antennal segment, no defects where observed in *Dll* expression in the head or in the thoracic primordia in *btd*^*ΔBTD-box*^/*Df(btd, Sp1*) embryos when compared with the wild type (Figure S1). These and previous results suggest that the BTD-box of *btd* and *Sp1* is not required for head segmentation neither ventral appendage specification.

The central nervous system (CNS) is generated from a small group of stem cells, NBs, which are specified during embryonic development [51]. Two types of NBs have been identified according to how they proliferate. Type I NBs divide asymmetrically, self-renewing and generating an intermediate cell, the ganglion mother cell (GMC), which divides only once to generate two cells that will differentiate as neurons or glial cells. Most of the brain NBs belongs to this type. Type II NBs also divide asymmetrically, self-renewing and generating an intermediate neural progenitor cell (INP). This INP will also divide asymmetrically several more times, self-renewing and generating GMCs that will divide only once. There are only eight type II NBs per hemibrain, but their progeny is much larger than that produced by type I NBs. Type I and II NBs can be easily distinguished because only type I NBs express the bHLH gene *asense* (*ase*) [52–54].

It has been suggested that Btd is also required to maintain type II NB identity along larval development. Thus, about 40% of the *btd* mutant clones ectopically express *ase* in type II NBs, which indicates that they are taking type I identity [19, 55].

Thus, we assessed whether the BTD-box plays a role in the maintenance of type II NB identity. To that end, as an indication of the type II NB transformation to type I NB, we looked at *ase* expression in type II NBs in third instar larval brains of animals in which the BTD-box was removed over the *Df(btd, Sp1*). To identify type II NBs, we used the genetic combination *worniu* (*wor*)-*Gal4 ase-Gal80*; *UAS-Cherry* (hereafter *wach*). *wor-Gal4*, a pan-neuroblast driver, will drive the expression of *UAS-Cherry* in all NBs, and *ase-Gal80* will repress the action of the Gal4 in type I NBs. The perdurance of the Cherry product also allows labeling most of the NB progeny as a Cherry-expressing cluster of cells. This *wach* genotype has been previously used successfully to identify type II NBs [20]. In *wach* wild type animals, we never observed expression of *ase* in type II NBs (Fig. 3A and B; n=37). In *btd*^*ΔBTD-box*^/*Df(btd, Sp1*); *wach* ganglia, we observed cases in which *ase* is expressed in type-II NBs (Fig. 3C; 20%, n=20). Although, the frequency in which this happens was lower than the reported frequency for expression of *ase* in *btd* null mutant clones [55]. No ectopic *ase* expression was observed in type II NBs of *Sp1*^*ΔBTD-box*^/*Df(btd, Sp1*) individuals (Fig. 3D; 0%, n=20), however in the double mutant *btd*^*ΔBTD-box*^, *Sp1*^*ΔBTD-box*^ or the *btd*^*ΔBTD-box*^, *Sp1*^*ΔBTD-box*^/*Df(btd,Sp1)* we observed 2% (n=67) and 30% (n=23) *ase* expression, respectively (Fig. 3E and F). Thus, we conclude that the BTD-box in Btd plays a role in the maintenance of type II NB identity.

**Figure 3:**
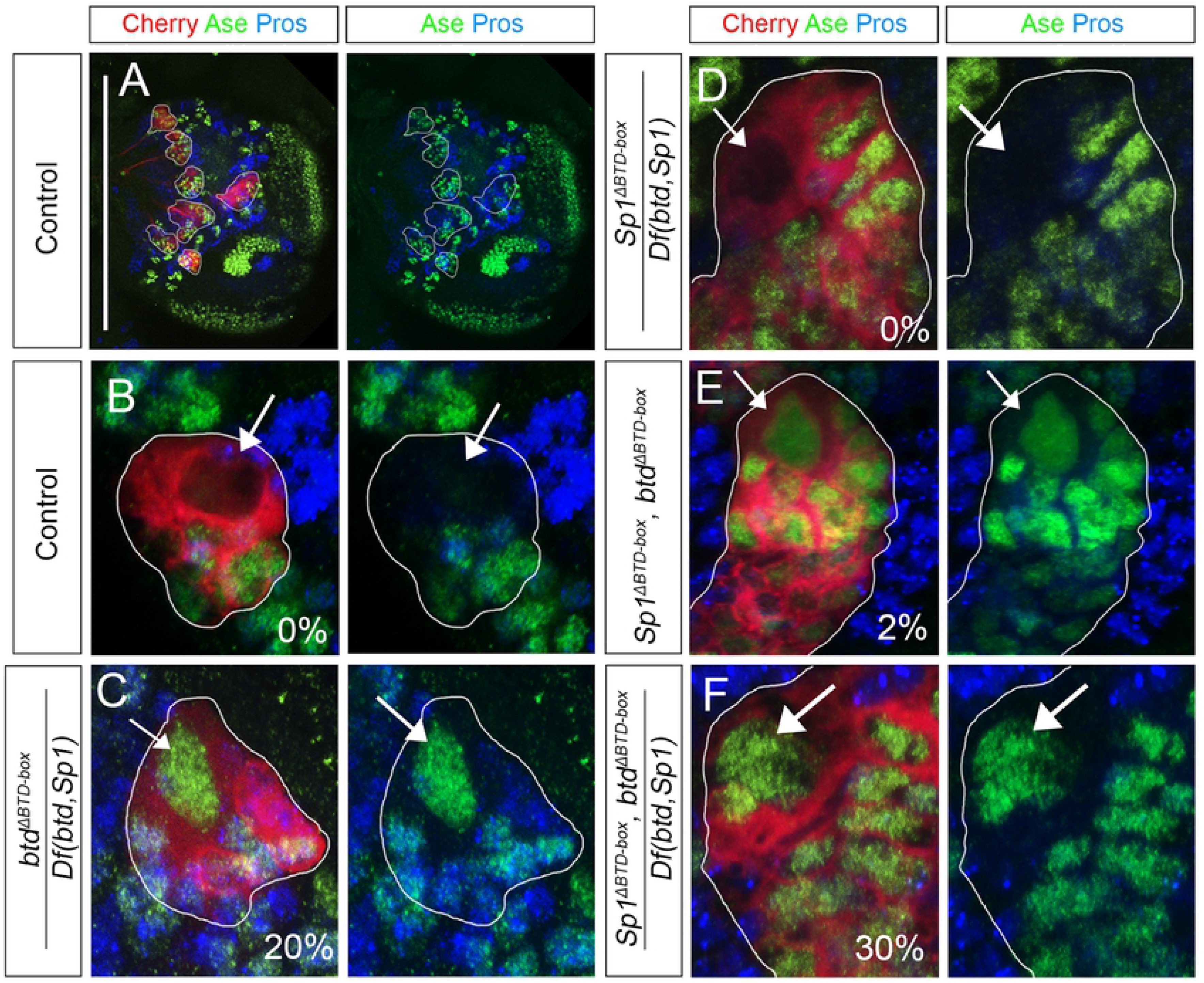
The BTD-box of Btd is required to maintain type II NB identity. (A-F) Stainings for Ase (green), Cherry (red) and Prospero (Pros, blue) in type II NB clusters of *wach* in third instar ganglia. (A) Dorsal view of a hemi-ganglia where seven of the eight *wach* clusters (red) can be observed (white outline). White bar indicates the midline. Anterior is up. (B) Close detail of *wach* clusters. Within the *cherry* expressing cells, type II NBs can be identified by their larger size and the lack of both nuclear Pros and Ase (white arrows). In *btd*^*ΔBTD-box*^/*Df(btd,Sp1); wach/+* (C) and *Sp1*^*ΔBTD-box*^, *btd*^*ΔBTD-box*^/*Df(btd,Sp1); wach/+* (F) ganglia Ase was present in some type II NB (white arrow). No derepression of *ase* was observed in type II NBs (white arrow) in *Sp1*^*ΔBTD-box*^/*Df(btd,Sp1); wach/+* (D) and *Sp1*^*ΔBTD-box*^, *btd*^*ΔBTD-box*^; *wach/+* (E) ganglia. Ase and Pros stainings are shown at the right for each panel. The % of *ase* expression in the *wach* NB is indicated for each experiment.

### Role of the BTD-box domain in *btd* and *Sp1* gain of function experiments

To further investigate the *in vivo* role of the BTD-box domain we have generated UAS versions of *Sp1* and *btd* where this domain was deleted, and tested their ability to reproduce the gain of function phenotypes of their counterpart wild type versions. As a control, we generated new *Sp1* and *btd* wild type UAS lines that were inserted in the same chromosomal location as the mutant UAS lines using ΦC31 mediated recombination to ensure similar expression levels (see Material and Methods)[56]. *btd* and *Sp1* have redundant functions regulating thoracic limb primordia *Dll* expression in late embryos, and only the elimination of both genes abolish *Dll* and *Dll-LT* CRM activation [13, 17]. The expression of mutant versions of *Sp1* and *btd* were able to rescue *Dll* and *Dll-LT* activity just as well as the control wild type versions when expressed in the second thoracic segment using the *prd-Gal4* line in a *Df(btd, Sp1)* mutant background (Figure Sup. 2). These and previous results suggests that the BTD-box is not required for the correct expression of *Dll* in the embryonic leg primordia.

We have previously shown that Sp1 is able to activate *Ser* expression in the tarsal segments of the leg [18]. Temporally restricted expression of *Sp1* with the *ptc*-*Gal4* line in the leg imaginal disc cause ectopic *Ser* expression in the tarsal segments with the consequent loss of *bib* (Figure 2I). This ectopic *Ser* activation by Sp1 is lost when the BTD-box deletion UAS version of *Sp1* is used instead of the wild type one (Figure 2J), which confirm our previous finding that *Sp1*^*BTD-box*^ impairs correct *Ser* activation in a sensitized background. To extend these results, we have also compared the phenotypes of the overexpression of *Sp1* and *btd* with and without the BTD-box domain in the leg. The expression of wild type versions of these genes with the *Dll*-*Gal4* driver truncates leg formation up to the distal tibia. However, these phenotypes were attenuated in the case of the BTD-box deletion versions, both causing less severe truncations than their wild type counterparts (Figure S3).

The ectopic expression of *Sp1* and *btd* in the wing imaginal disc, where they are not normally expressed, activated the leg patterning genes *Dll* and *dac* and caused wing-to-leg transformations in the adult appendage [13, 17] (Figure 4A, B and D). These phenotypes are weaker when the mutated versions, *Sp1*^*ΔBTD-box*^ or *btd*^*ΔBTD-box*^ were used instead of the wild type ones. Both, the ability to induce *Dll* and *dac* expression and the wing-to-leg transformations were reduced in ΔBTD-box mutant versions (Figure 4C and E). These differences are more evident in the case of *Sp1* gain of function experiments where the overgrowths induced by the wild type *Sp1* and the ectopic activation of *Dll* and *dac*, with the consequent transformation of the wing tissue to leg fates, were clearly reduced in the *Sp1*^*ΔBTD-box*^ mutant version (compare 4B with 4C). In summary, we conclude that the BTD-box domain in Sp1 and Btd is not essential for leg development. However, mutant flies that lack this domain in a sensitized genetic background (*Df(btd, Sp1)*) or the ectopic expression experiments of the BTD-box deletion lines, reveal a role for the BTD-Box in the transcriptional output of Sp1 and Btd.

**Figure 4:**
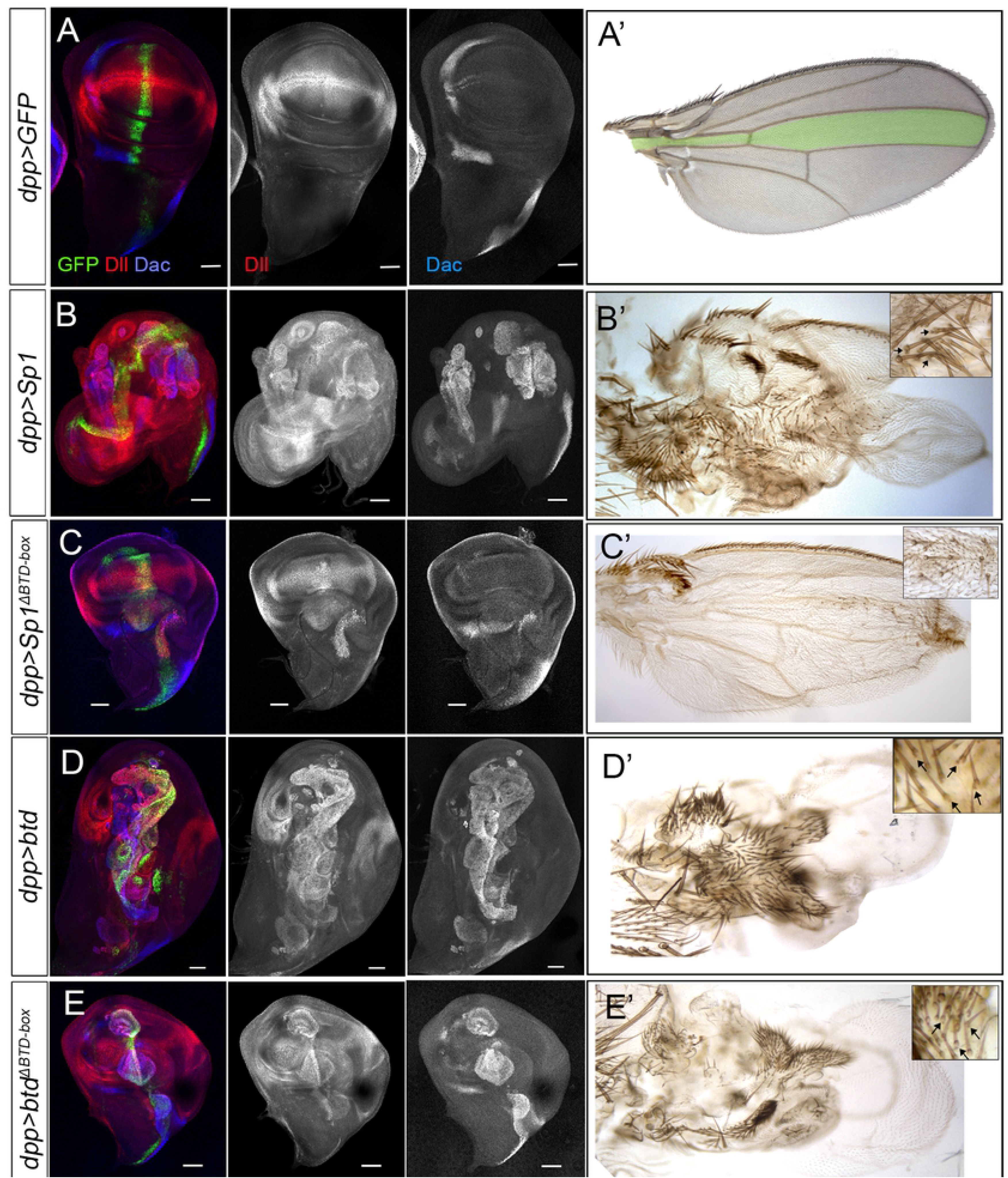
Ectopic expression of wild type and BTD-box deletion versions of *Sp1* and *btd* in the wing disc. (A-E) Ectopic expression of UAS-*GFP* (A), UAS-*Sp1* (B), UAS-*Sp1*^*ΔBTD-box*^ (C), UAS-*btd* (D) and UAS-*btd*^*ΔBTD-box*^ (E), under the control of *dpp-Gal4* in the wing imaginal disc and the corresponding adult wing at each side (A′-E′). Imaginal discs were stained for *dpp-Gal4* expression (green), Dll (red), Dac (blue). Insets show a high magnification of the ectopic leg tissue with the characteristic associated bract, a typical feature of leg bristles (black arrows). Note that the ectopic expression of *Sp1* (B) strongly activates *Dll* and *dac* expression and cause a dramatic overgrowth of the tissue causing a wing-to-leg fate transformation as observed by the presence of bracted bristles characteristic of leg tissue. These phenotypes are attenuated when the *Sp1*^*ΔBTD-box*^ (C) version is used instead of the wild type. Note the presence of some ectopic bristles without bracts in the wing (C′). The ectopic expression of *btd* (D) also activates *Dll* and *dac* and induced wing to leg transformation (D′). Less severe activation of *Dll* and *dac* and leg transformation was observed when the BTD-box deletion version of *btd* was used in place of the wild type (E′).

### Sp1 represses antennal fate genes in the leg

In a previous transcriptomic analysis of *Sp1* mutant leg discs, the antennal specific genes *danr* and *ss* were identified as potential targets of Sp1 [18]. Accordingly, in flies where two copies of *Sp1* and one of *btd* were mutated, some legs presented distal antenna-like structures similar to the arista, suggesting a role for *Sp1* (and possibly *btd*) repressing antennal fates [18]. To investigate this new role in more detail, we have analyzed the expression of the antennal-determinant genes *hth*, *sal*, *ss* and *danr* in mutant clones for *Df(btd, Sp1)* generated in second instar leg disc (48-72 hrs AEL). To monitor *danr* expression we have identified a dedicated CRM (named *danr1-GFP*) that reproduces its expression in the antenna and, just as the endogenous *dan/danr* genes, it is not activated in the leg disc (See Material and Methods and Figure S4) [47, 48]. Interestingly, only *ss* and *danr* are derepressed in *Df(btd, Sp1)* mutant clones, and only in those clones located in the distal domain of the leg (Figure 5A, B and E). Next, we made single *btd*^*XG*^ and *Sp1*^*HR*^ null mutant clones to determine whether both genes are required for *ss* and *danr1-GFP* repression. Only in *Sp1*^*HR*^ mutant clones we observed *ss* and *danr1-GFP* expression, demonstrating that *Sp1* is required to repress these antennal genes in the leg imaginal disc (Figure 5C, D, F and G).

**Figure 5:**
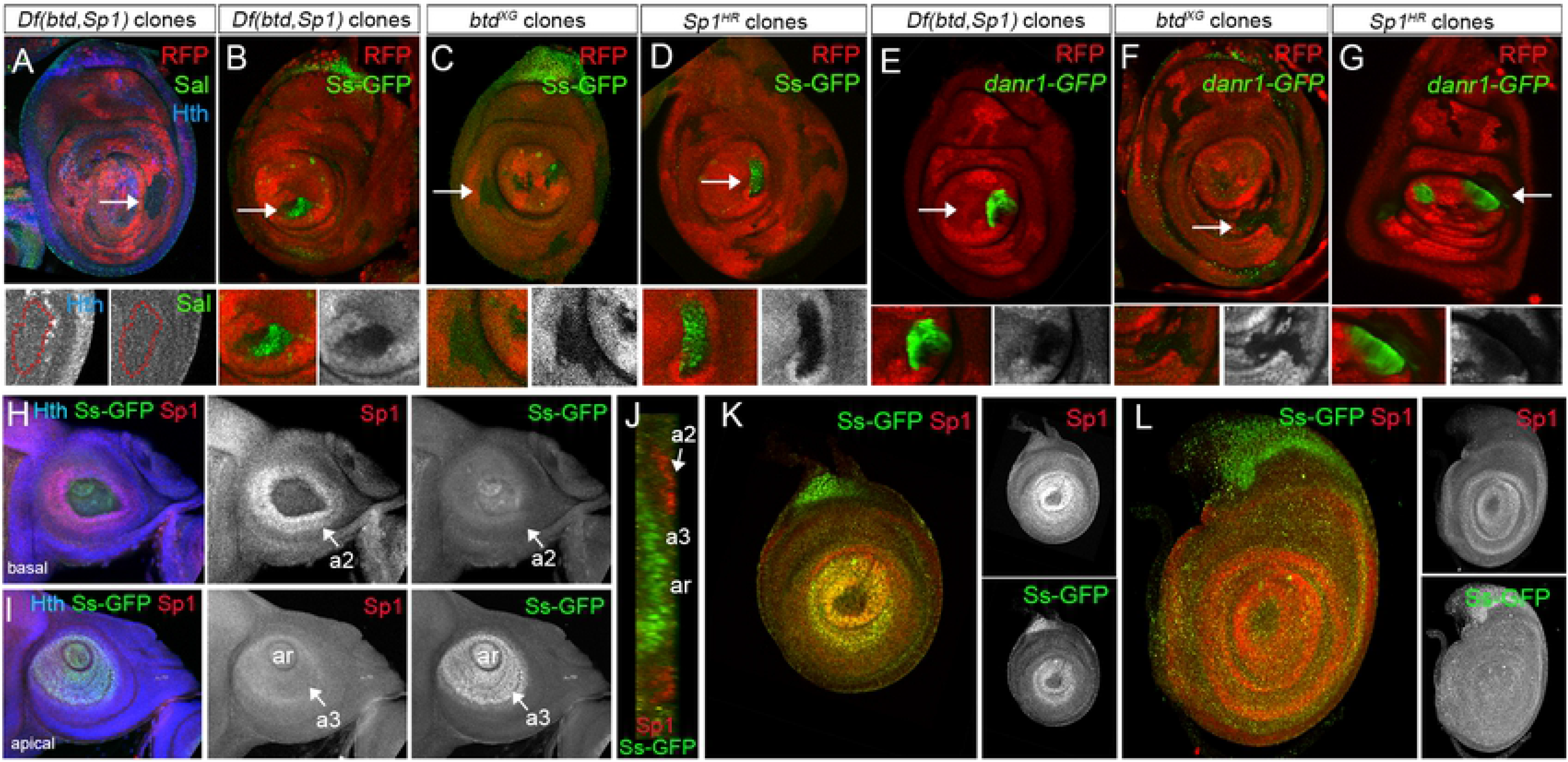
Sp1 represses *ss* and *danr1-GFP* expression in the leg. (A-G) Third instar leg imaginal discs presenting clones generated 48-72 hrs AEL for *Df(btd,Sp1)* (A, B and E), *btd*^*XG81*^ (C and F) and *Sp1*^*HR*^ (D and G) marked by the absence of RFP (red). Below each panel a close up of a clone is shown. (A-B) Leg discs with *Df(btd,Sp1)* mutant clones stained for Sal (green) and Hth (blue) in A and for Ss-GFP (green) in B. Note the derepression of *ss-GFP* within the clone (arrow). (C-D) Leg discs with *btd*^*XG81*^ (C) and *Sp1*^*HR*^ (D) mutant clones stained for Ss-GFp. Note the derepression of *ss-GFP* only in *Sp1*^*HR*^ mutant clones (arrows). (E-G) Leg imaginal discs with *Df(btd,Sp1)* (E), *btd*^*XG81*^ (F) and *Sp1*^*HR*^ (G) mutant clones and stained for *danr1-GFP*. Note the derepression of *danr1-GFP* only in *Df(btd,Sp1)* and *Sp1*^*HR*^ mutant clones (arrows). (H-J) Basal (H), apical (I) and cross-section (J) views of an antenna imaginal disc stained for Hth (blue), Ss-GFP (green) and Sp1 (red). Separate channels are also shown. a2: second antennal segment; a3: third antennal segment; ar, arista. Note that Ss-GFP and Sp1 are present in non-overlapping antennal segments. (K and L) Early third (K) and late third (L) instar leg imaginal discs stained for Sp1 (red) and Ss-GFP (green). Single channels are shown next to each panel. Note that *Sp1* and *ss-GFP* are coexpressed in early third instar discs but not in mature discs.

As *ss* is expressed in the antenna and transiently in the leg, we decided to compare the spatial and temporal dynamics of *Sp1* expression with that of *ss* in the leg and antenna imaginal discs. In the antenna, *Sp1* is expressed in a single ring in the second segment (a2) while *ss* and *dan/danr* are restricted to the third (a3) and arista segments (Figure 5H-J and Figure S4) [46, 47]. In the leg disc, *ss* is transiently expressed from second to early third instar in a broad ring in the distal domain, that corresponds to the future tarsal segments of the leg [46]. This early *ss* expression overlaps with Sp1, which extends more medially (Figure 5K). In late third instar leg imaginal discs, *ss* expression decays while Sp1 continues to be expressed (Figure 5L).

As it has been proposed that *dan/danr* act downstream of Ss in the antenna [47], it is reasonable to suggest that Sp1 repression of *danr1-GFP* could be indirect through the regulation of *ss*. To test this possibility, we first identified and mutated all the Sp1 putative binding sites in the *danr1*-CRM (eight in total) with almost no effect on its activity. *danr1*^*Sp1 Mut*^-*GFP*, as the wild type version (*danr1-GFP*), is normally activated in the antenna but not in the leg discs, at the exception of few scattered cells in the leg (Figure S4). Next, we tested whether the *danr1-GFP* derepression observed in *Sp1* loss of function conditions depends on Ss. To this end, we first confirmed that ectopic expression of *ss* in the leg activated *danr1*-*GFP* (Figure 6A and B) and that *danr1*-*GFP* activity is derepressed in *Sp1*^*HR*^ mutant leg disc in the distal domain (compare Figure 6C with 6A). Next, we analyzed *danr1*-*GFP* activity in *Sp1*^*HR*^ mutant leg imaginal discs that simultaneously downregulated Ss levels in the anterior compartment (*Sp1*^*HR*^; *ci>ss*-RNAi). Knockdown of Ss in a *Sp1*^*HR*^ mutant maintain *danr1*-*GFP* repressed, suggesting that *danr* expression is activated by Ss, and is thus indirectly repressed by Sp1 (compare Figure 6D with 6C). Accordingly, a recent genome wide *in vivo* profile of Sp1 and Dll binding by chromatin immunoprecipitation (ChIP) has shown that these transcription factors bind close to the transcription start of *ss* [31].

**Figure 6:**
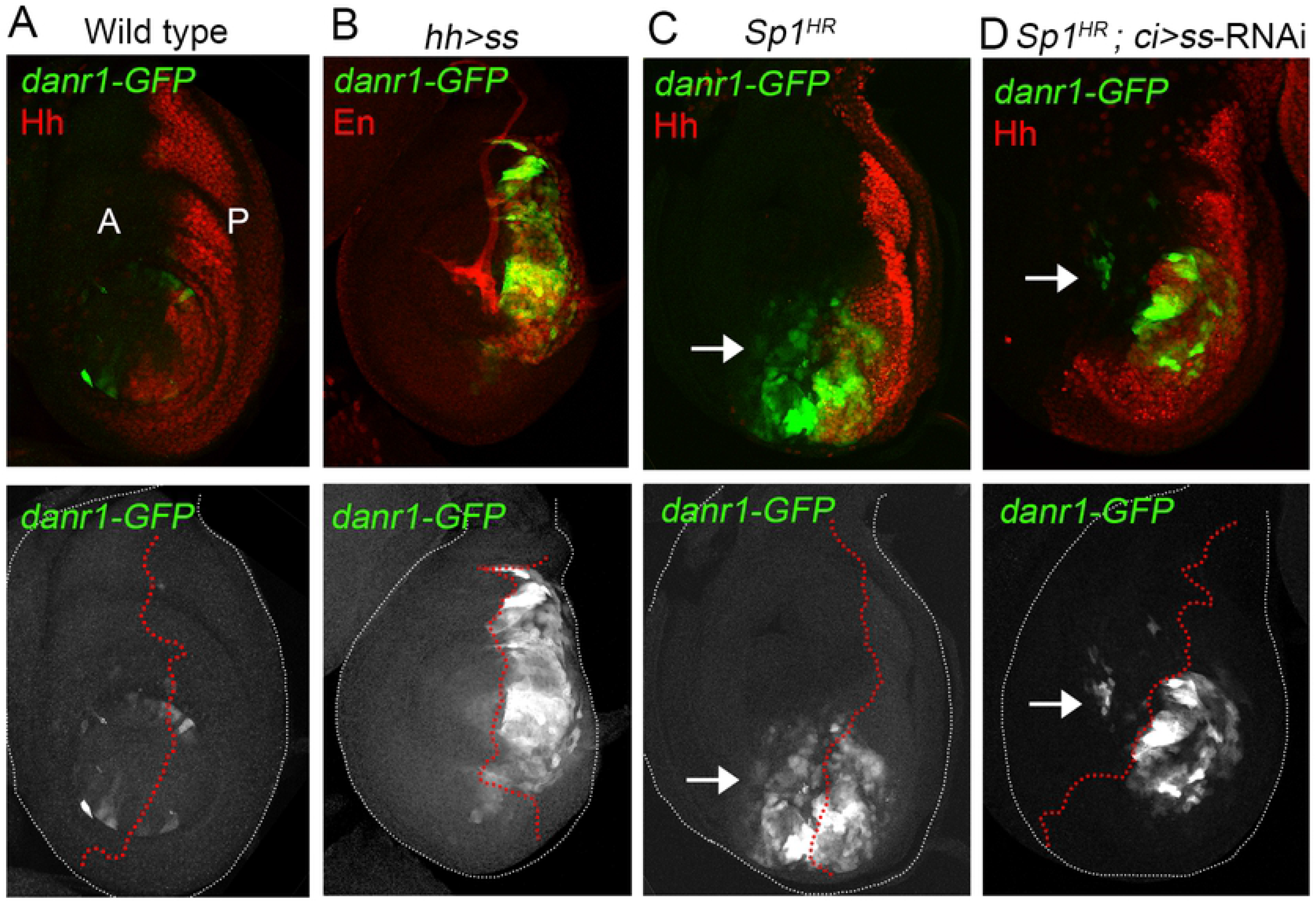
Sp1 represses *danr1-GFP* activity through Ss. (A-D) Third instar leg imaginal discs stained for *danr1-GFP* (green) and *hh-dsRED* (red) or En (red) in B. Below each panel the *danr1-GFP* channel is shown and the antero-posterior compartment boundary is depicted with red dots. A, anterior; P, posterior. (A) Wild type control disc. (B) *hh-Gal4*, UAS-*ss* activates *danr-1-GFP* expression in the posterior compartment. (C) In a *Sp1*^*HR*^ mutant leg disc, *danr1-GFP* is derepressed in the distal domain in both compartments. (D) In a *Sp1*^*HR*^ mutant leg disc, the knockdown of Ss in the anterior compartment prevents the derepression of *danr1-GFP* when compared with the posterior compartment. Note that few cells still express *danr1-GFP* in the anterior compartment (arrow).

### The BTD-Box domain of Sp1 is required for proper *ss* and *danr1*-CRM repression

We have shown that *ss*, and therefore *dan/danr* are targets of Sp1 in the leg. We decided to use both genes as readouts of Sp1 transcriptional activity in *Sp1* mutant versions that lack the BTD-box. Consistently with our clonal analysis, *ss* and *danr1-GFP* are strongly derepressed in a distal ring in late third instar *Sp1*^*HR*^ mutant leg discs (Figure 7B and G). Although *ss* derepression is not observed in *Sp1*^*ΔBTD-box*^ mutant leg discs, a few cells show ectopic activation of the *danr1-*CRM in this mutant condition (Figure 7C and H). Remarkably, in the sensitized background (*Sp1*^*ΔBTD-box*^/*Df(btd, Sp1)* a strong derepression of *ss* and *danr1-GFP* activity could be observed in mature third instar leg discs (Figure 7D and I). This derepression was never observed in the control *Sp1*^*Wt*^/*Df(btd, Sp1)* animals (Figure 7E and J).

**Figure 7:**
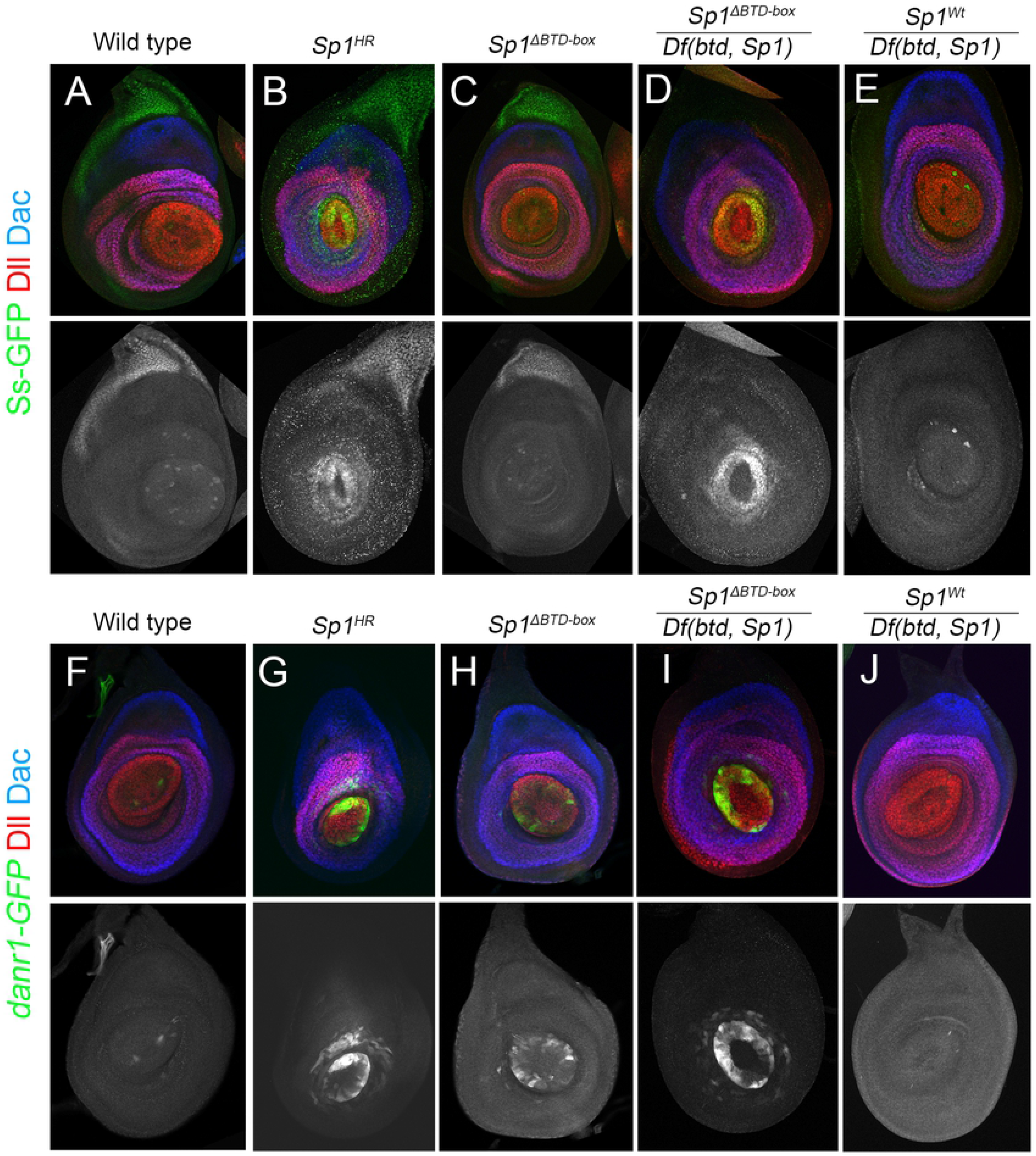
The BTD-box of Sp1 is required for the correct repression of *ss* and *danr1-GFP* expression in the leg. (A-E) Late third instar leg imaginal discs stained for Dll (red), Dac (blue) and Ss-GFP (green) of wild type (A), *Sp1*^*HR*^ (B), *Sp1*^*ΔBTD-box*^ (C), *Sp1*^*ΔBTD-box*^/*Df(btd,Sp1)* (D) and *Sp1*^*Wt*^/*Df(btd,Sp1)* (E) mutant animals. Below each panel the single channel for Ss-GFP is shown. (F-J) Mature third instar leg imaginal discs stained for Dll (red), Dac (blue) and *danr1-GFP* (green) of wild type (F), *Sp1*^*HR*^ (G), *Sp1*^*ΔBTD-box*^ (H), *Sp1*^*ΔBTD-box*^/*Df(btd,Sp1)* (I), and *Sp1*^*Wt*^/*Df(btd,Sp1)* (J) mutant animals. Below each panel the single channel for *danr1-GFP* is shown.

Temporally restricted expression of BTD-box deficient versions of *Sp1* and *Btd* in the antenna with the *hh-Gal4, Gal80*^*ts*^ system also supports the idea that the BTD-box is required for the efficient repression of *ss*. While the ectopic expression in the antenna of *Sp1* and *btd* strongly repressed *ss* expression, this effect is less dramatic when missexpressing the BTD-box deficient versions (Figure 8).

**Figure 8:**
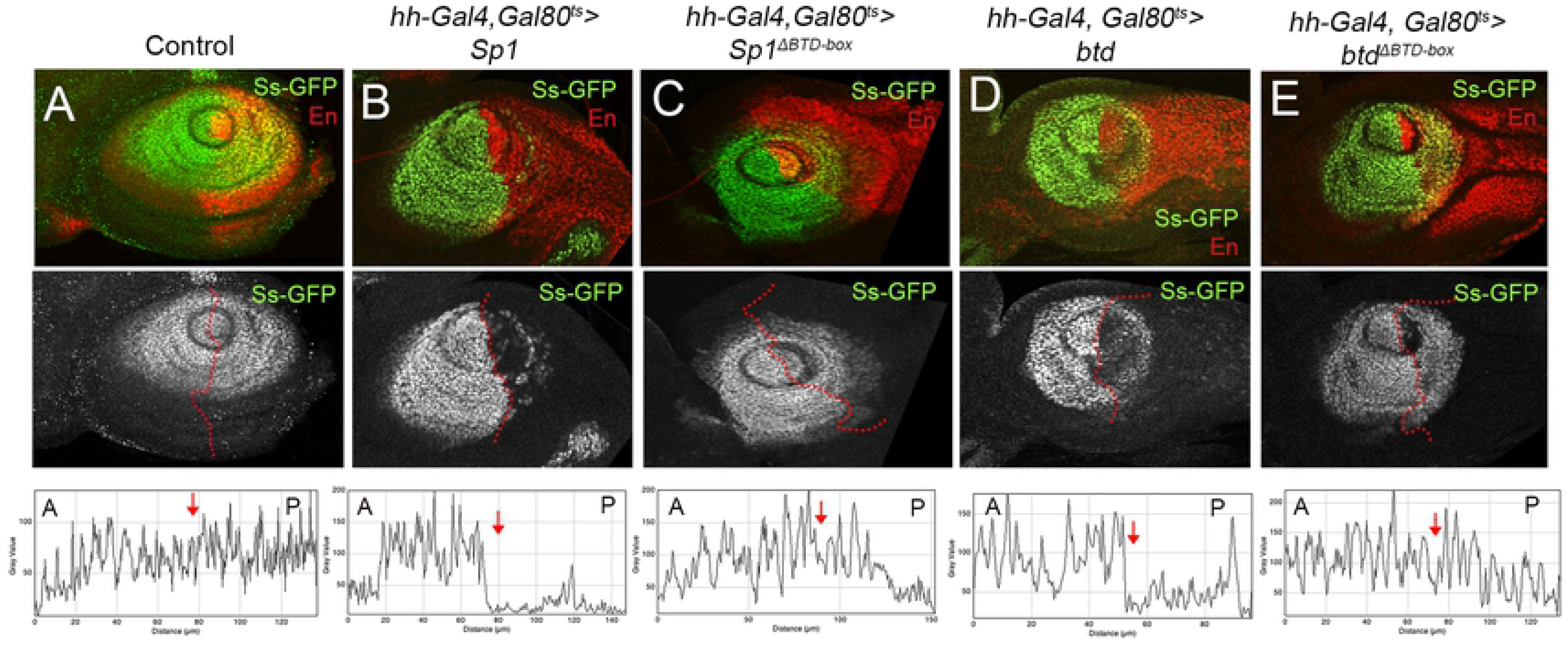
The BTD-box of Sp1 and Btd is required for the correct repression of *ss* in gain of function experiments in the antenna. (A-E) Third instar antenna imaginal discs stained for Ss-GFP (green) and En (red) where either no UAS (A), UAS-*Sp1* (B), UAS-*Sp1*^*ΔBTD-box*^ (C), UAS-*btd* (D) or UAS-*btd* ^*ΔBTD-box*^ (E) were ectopically expressed starting from third instar larval stage using the *hh-Gal4*; *tubGal80*^*ts*^ driver. Below each panel the single channel for Ss-GFP is shown and the antero-posterior boundary is depicted in red. Ss-GFP intensity plots of the corresponding images shown in A to E are also shown. Red arrow marks the antero-posterior boundary. A, anterior; P, posterior. All larvae were dissected at the same time and a representative image is shown for each experiment.

All together, these results confirm the role of the BTD-box domain of Sp1 for the effective repression of the antennal gene *ss*.

## Discussion

In this work we have analyzed the *in vivo* requirements for the conserved BTD-box domain of the Sp family of transcription factors in *Drosophila*. Using gene-editing tools we have generated new *Sp1* and *btd* alleles and UAS versions that lack this domain. In addition, we have identified a new role for Sp1 as a repressor of *ss* expression and distal antennal fates in the leg, a function that partly requires the presence of the BTD-box domain.

In *Drosophila*, the members of the Sp family *Sp1* and *btd*, have both unique and partially redundant functions throughout development. During early embryogenesis, *btd* contributes to head segmentation and later on *btd* and *Sp1* act redundantly to activate *Dll* expression in thoracic ventral limb (leg) primordia in late embryos and in leg imaginal discs [13, 17, 49]. Later in larval development, these genes have divergent contributions to leg growth [17, 18]. In the nervous system, both Btd and Sp1 are required for type II NB specification and later on *btd* prevents the premature differentiation of intermediate neural progenitor cells [19, 20, 55]. Unexpectedly, we found that animals that lack the BTD-box of *Sp1* or *btd* are viable, fertile and have nearly wild type legs. However, the viability of the double *Sp1*^*ΔBTD-box*^, *btd* ^*ΔBTD-box*^ mutant is strongly compromised as few animals where found that pass larval development. Importantly, when confronting the *Sp1*^*ΔBTD-box*^ or *btd* ^*ΔBTD-box*^ alleles with a deficiency for both genes, thus reducing their gene dosage, the phenotypes are more dramatic. Specifically, for *Sp1*^*ΔBTD-box*^/*Df(btd, Sp1)* we observed a clear reduction on leg size and defects on leg joint formation, phenotypes resembling those described for *Sp1* null mutants [17, 18]. Analysis of leg imaginal discs of this genotype revealed defects on the correct regulation of *ss* and *Ser* expression.

Moreover, we found that type II NB identity maintenance was compromised in the sensitized background for the *btd* ^*ΔBTD-box*^ allele. The fact that we can observe clusters of *cherry*-expressing cells including a NB that expresses *ase* indicates that, at any time, during larval development, the NB has lost its type II identity. The low frequency of the event suggests an erratic behavior that, on the other hand, we do not know if it is reversible, but that obviously indicates an implication of the BTD-box in the maintenance of their identity during development.

These results suggested that although the BTD-box of Sp1 or Btd is not essential during development, in a sensitized background a requirement of this domain to correctly regulate its target genes is revealed. Importantly, the function of the BTD-box seems to be target specific. For example, embryonic head formation and appendage specification do not rely on the presence of the BTD-box of Sp1 or Btd. However, activation of *Ser* and repression of *ss* and *danr* in the leg and type II NB identity maintenance in the brain were affected in a sensitized background for *Sp1* and *btd* BTD-box mutant alleles.

Aligned with our results, *in vitro* analysis of the transcriptional activity of the human Sp1 transcription factor revealed that a deletion that contained the BTD-box was not essential for the regulation of all Sp1 target genes [6, 7, 57, 58]. However, the BTD-box was shown to be critical for efficient activation of the low-density lipoprotein (LDL) receptor promoter [7, 8, 58]. This promoter requires the cooperative binding of the sterol regulatory element binding proteins (SREBPs) and of Sp1. Importantly, deletion of the BTD-box domain abolished the stimulated binding of Sp1 by SREBP in this promoter [8]. Therefore, the Sp1 BTD-box could function as a transactivation domain required to stimulate effective binding of Sp1 and its cofactors to their target genes. Our analysis of several *Drosophila* Sp1 target genes suggests a similar function for the BTD-box domain. Previously, we have proposed that Sp1 interacts with PD transcription factors, such as Ap, to activate Ser expression in the presumptive tarsal joints [18]. Our results suggested a requirement for the BTD-box of Sp1 for effective *Ser* regulation in loss and gain of function experiments. In addition, we propose a role for the BTD-box of Btd in type II NB maintenance where it has been demonstrated that the transcriptional effector of the EGFR pathway, PntP1, function cooperatively with Btd [19, 55].

In this work we have also identified *ss* as a novel Sp1 target gene. *ss* is transiently expressed in the leg disc where it is required for tarsal development while in the antenna specifies distal structures. Importantly, forced expression of *ss* in the leg induced the formation of distal antenna structures such as the arista [46]. We have shown that Sp1 represses *ss* expression in the leg and the antenna, however in early third instar leg imaginal discs both genes are coexpressed. Therefore, we propose that Sp1, in combination with an unidentified factor, turn down *ss* expression in the mature third instar leg discs. This repression is essential to prevent the activation of genes required for distal antenna specification, such as *dan/danr* that are activated by Ss [47]. The interaction between Sp1 and a hypothetical cooperative transcription factor could be mediated by the BTD-box, as the deletion of this domain prevents the correct repression of *ss* in gain of function experiments and in the *Sp1*^*ΔBTD-box*^/*Df(btd, Sp1)* mutant background.

In summary, our results highlight a role for the BTD-box for the precise regulation of several Sp1 and Btd target genes. Nevertheless, mutant animals that lack this domain efficiently regulate these genes and have nearly normal appendages. To explain the great degree of conservation of the BTD-box domain, it is possible to imagine that its roles have been gradually reduced during evolution or selected for specific subsets of target genes.

## Materials and Methods

### Generation of the Sp1^ΔBTD-box^ and btd^ΔBTD-box^ mutant alleles

To generate the *Sp1*^*ΔBTD-box*^ mutant flies we used the *Sp1*^*HR*^ mutant allele, which has an *attP* integration site in place of the third exon of Sp1 generated by homologous recombination, to insert a version of this exon without the BTD-box [59]. The wild type version of the third exon of *Sp1* and the version lacking the BTD-box were cloned in the reintegration vector (RIV) using the following primers:

5′-cagt**gctagc**gcagctggccctatacgactcgcg-3′ (**NheI**) and 5′-cagt**ctcgag**cgctgttttagttggtgaacaggaagg-3′ (**Xho**)

To generate the deletion of the BTD-box within the third exon of *Sp1* we replaced the 30 bp sequence encoding for the BTD-box for a StuI site that maintains the reading frame using these primers:

5′-cagt**aggcct**ggcataacgcctctgggatctg-3′and 5′-cagt**aggcct**caggaggcggaacgcctgggtc-3′

*Sp1*^*Wt*^ and *Sp1*^*ΔBTD-box*^ RIV vectors were then injected in *Sp1*^*HR*^/*Dp*(*1;Y*)*lz*+ mutant embryos. Candidate transformants were subjected to CRE mediated excision of the reporter gene and selectable marker and the results were confirmed by sequencing.

The *btd*^*ΔBTD-box*^ allele was generated by CRISPR/Cas9, using pCFD3 vector for driving gRNA expression and a germline-expressing Cas9 donor strain in a wild type and in a *Sp1*^*ΔBTD-box*^ background. The following sequence was used as gRNA: 5′-gtcgggcacgtgcagcgaacggag-3′. Candidate mutant alleles were sequenced and selected only those that delete 4 or more aa of the BTD-Box domain, while maintaining the reading frame.

### Generation of UAS-Sp1, UAS-Sp1 ^ΔBTD-box^, UAS-btd and UAS-btd ^ΔBTD-box^

Wild type *Sp1* and *btd* cDNA was cloned in a pUAST attB vector as described in [17]. To generate the *Sp* ^*ΔBTD-box*^ and the *btd* ^*ΔBTD-box*^ we used the QuickChange Site-Directed Mutagenesis Kit (Stratagene) to delete the 30 bp encoding for the BTD-box using the following primers:

For *Sp1*^*ΔBTD-box*^ 5′-agatcccagaggcgttatgcccaggaggcggaacgcctgggtccc-3′ and 5′-gggacccaggcgttccgcctcctgggcataacgcctctgggatct-3′
For *btd*^*ΔBTD-box*^ 5′-tccagacgcaattgcaacgcaccaacgagatgagcggcctg-3′and 5′-caggccgctcatctcgttggtgcgttgcaattgcgtctgga-3′

To ensure similar expression levels, all UAS constructs we inserted into the same attP site (86Fb).

### Identification of the danr1 CRM and Sp1 binding site mutagenesis

Using the Fly Light, Fly Enhancers databases and self-made constructs, we screened the *dan* and *danr* genomic region for DNA fragments that drives expression of the *Gal4* gene in a *dan/danr*-like expression pattern [60, 61] (Figure S4). We selected *dan1*, *danr1* and *danr2* fragments as they presented open chromatin profiles by FAIRE seq that were specific for the eye-antenna disc and remain closed in the leg disc [62](Figure S4). DNA fragments were cloned first into the pEntry/D-TOPO vector and then swapped into the attB-pHPdesteGFP vector, replacing the *ccdB* gene with the enhancer sequence using the LR Clonase Enzyme Mix (ThermoFiser) [63].

The following primers were used:

*dan1 sense:* 5′-**cacc**ctccattgcattgcattgcattg-3′ *dan1asense:* 5′-gtatgtatgtaggtaaccaggg-3 *danr1 sense:* 5′-**cacc**gggttttcaattacagcggttag-3′ *danr1 asense:* 5′-ttttgctggccaaattggtgccag-3′
*danr2 sense:* 5′-**cacc**gtctttctcgcccattttcgc-3′ *danr2 asense:* 5′-accgtgtcttcctagcccaccg-3′

From all these fragments, only one element named *danr1*-CRM faithfully reproduced *dan/danr* expression pattern in the antenna and leg imaginal discs.

Eight potential Sp1 binding sites were identified in *danr1* CRM on the basis of a bioinformatic analysis from the JASPAR CORE Insecta database (http://jaspar.genereg.net/). Mutagenesis of the putative Sp1 binding sites was performed using the QuikChange Site-Directed Mutagenesis Kit (Stratagene). All the reporter constructs were inserted and analyzed at the same landing attP site. The sequence of all primers used in this study:

*danr1-Sp1 binding site 1 sense*
5′-aggagcggcccaagaagcagccttctattcaagagcagccgccacggatg-3′
*danr1-Sp1 binding site 1 asense*:
5′-catccgtggcggctgctcttgaatagaaggctgcttcttgggccgctcct-3′
*danr1-Sp1 binding site 2 sense*
5-tccgcaccggtggccgatggagttttatggacatgaaaatggcgatgaatt-3′
*danr1-Sp1 binding site 2 asense*
5′-aattcatcgccattttcatgtccataaaactccatcggccaccggtgcgga-3′
*danr1-Sp1 binding site 3 sense*
5′-agctcgttcgacatttttggactcttatgtgttaatgtaaggataatcac-3′
*danr1-Sp1 binding site 3 asense*
5′-gtgattatccttacattaacacataagagtccaaaaatgtcgaacgagct-3′
*danr1-Sp1 binding site 4 sense*
5′-gcgcatatgcatttttgattaaataaacacgtccatgtagctcgaagtca-3′
*danr1-Sp1 binding site 4 asense*
5′-tgacttcgagctacatggacgtgtttatttaatcaaaaatgcatatgcgc-3′
*danr1-Sp1 binding site 5 sense*
5′-tggcattctggttttggttcgtttatcttcttaaattactccagccaatc-3′
*danr1-Sp1 binding site 5 asense*
5′-gattggctggagtaatttaagaagataaacgaaccaaaaccagaatgcca-3′
*danr1-Sp1 binding site 6 sense*
5′-atggcaccatggccagcttctattaccaagaagcagccttctatt-3′
*danr1-Sp1 binding site 6 asense*
5′-aatagaaggctgcttcttggtaatagaagctggccatggtgccat-3′
*danr1-Sp1 binding site 7 sense*
5′-ctgaaaagttcaatgccaaaaatcgcccaaagtgacgcgcagaat-3′
*danr1-Sp1 binding site 7 asense*
5′-attctgcgcgtcactttgggcgatttttggcattgaacttttcag-3′
*danr1-Sp1 binding site 8 sense*
5′-ctccagtctcggcacgaatcgattaagaagcagagctggtggatc-3′
*danr1-Sp1 binding site 8 asense*
5′-gatccaccagctctgcttcttaatcgattcgtgccgagactggag-3′

### Drosophila strains

*Sp1*^*HR*^ is a null mutant allele where the third exon that contains the DNA-binding domain and the BTD-box is replaced with an attP integration site using homologous recombination [18]. *btd*^*XG81*^ is a null allele and the *Df(btd,Sp1)* deletes *btd*, *Sp1*, and two adjacent genes with unknown function (CG1354 and CG32698) [5, 17]. UAS-*btd RNAi, Dll-LT-lacZ, Dllm-Gal4 (Dll-Gal4*^*212*^, *UAS-flp; act-FRT- stop-FRT-Gal4, UAS-GFP), bib-lacZ* and a duplication on the Y chromosome that covers the *btd* and *Sp1* genes (*Dp(1;Y)lz+)* have been described previously [17, 18, 38]. *The Dll-Gal4*^*212*^, *dpp-Gal4, ptc-Gal4, tubGal80ts, hh-Gal4, hh-dsred, Ci-Gal4*, UAS-*ss-RNAi* and the *ss-GFP* line (BL: 42289) are all available at Bloomington Stock Center.

For loss-of-function clonal analysis we used the following genotypes: *yw btd*^*XG81*^, *Sp1*^*HR*^ or *Df(btd,Sp1) FRT19A*/ *hsflp ubi-dsred FRT19A.* Larvae were heat shocked for 1 h at 37°C 48-72 hrs after egg laying (AEL). Temporally restricted gain-of-function experiments were performed using the *ptc-Gal4; tubGal80ts* and *hh-Gal4; tubGal80ts* system, which allowed temporal expression restriction of the different UAS lines to mid-third instar stage. *ptc-Gal4 or hh-Gal4; tubGal80ts* flies were crossed with each UAS strain, and the eggs laid each 24 h were collected and maintained at restrictive temperature (17°C) for seven days, when the fly vials were shifted to the permissive temperature (29°C) for two days before dissection.

For the analysis of the role of the BTD-box in the maintenance of type II NBs identity, *Dp(1;Y)lz+/Df(btd Sp1); wach/CyO, act-Gal4*, UAS-*GFP* males were crossed with: 1) *Sp1*^*ΔBTD-box*^, 2) *btd*^*ΔBTD-box*^ or 3) *Sp* ^*ΔBTD-box*^, *btd* ^*ΔBTD-box*^/*FM7, act-Gal4*, UAS-*GFP* females. Also, *wach* males were crossed with *Sp* ^*ΔBTD-box*^, *btd* ^*ΔBTD-box*^/*FM7, act-Gal4*, UAS*-GFP* females. Then, third instar larva males of the corresponding phenotypes were selected, dissected and stained with rabbit anti-Ase and mouse anti-Pros. *wach*: *wor-Gal4 ase-Gal80; 20x*UAS-*6x*UAS-*Cherry:HA* [20]

### Immunostaining

Embryos and larval and prepupal leg discs were stained following standard procedures [13]. Primary antibodies used were: rabbit and mouse anti-βGal (1/1000; Promega and MP Biomedicals, respectiveley), rat anti-Ser (1/1000; a gift from Ken Irvine, Rutgers University), rat anti-Sp1 (1/50; kindly provided by Richard Mann, Columbia University), rabbit anti-Ase (1:500 [20]), guinea pig anti-Dll and rabbit or guinea pig anti-Hth (1/2000; [38]), mouse anti-Pros (1/50; DSHB #MR1A), rabbit anti-GFP (1/1000; Thermo Fisher), mouse anti-Dac (1/50; DSHB #mAbdac1-1), mouse anti-En (1/50; DSHB #4D9), mouse anti-Sxl (1/100; DSHB #M18) and rabbit anti-Sal (1/200; a gift from Jose Felix de Celis). We used anti-Sxl to label all the somatic female nuclei to discriminate male from female embryos.

## Acknowledgements

We thank Richard S. Mann, Ken Irvine, Luis Alberto Baena, the Bloomington Stock Center, the Vienna Drosophila Resource Center and the Developmental Studies Hybridoma Bank at The University of Iowa for fly stocks and reagents. We specially thank the Confocal microscopy service and Eva Caminero and Mar Casado for fly injections.

---

**Figure S1: *Dll* expression is not altered in *Sp1* and *btd* BTD-box deletion mutants.** (A-C) Stage 13 embryos stained for Dll (green) and En (red) in A and B and Sxl (red) in C and D. an, antennal segment; mx, maxilar segment; lb, labial; T1-T3, thoracic segments. (A) Wild type embryo where the expression of *Dll* can be observed in the head segments and the ventral thoracic primordia. A white arrow marks the antennal segment. (B) *btd*^*XG*^ mutant embryo. Note the absence of the antennal segment (white arrow) and the reduction of *Dll* expression in the thoracic primordia. (C) *btd*^*ΔBTD-box*^/*Df(btd,Sp1)* mutant embryo. No defects on *Dll* expression in the head or the thoracic primordia are observed. Sxl staining was used to identify female embryos.

**Figure S2: Activation of *Dll* and *Dll-LT* CRM do not require the BTD box of Sp1 or BTD.** (A) Cartoon of a stage 14 embryo where the expression of *Dll* (green) and the activity of the *Dll-LT* CRM (red) in the thoracic segments is depicted. (B and C) Wild type embryo (B) and *Df(btd,Sp1)* (C) stage 14 mutant embryo stained for Dll (green) and *Dll-LT*-*lacZ* (red). Below the panels, the single channel for *Dll-LT* is shown. *Dll* expression is greatly reduced and LT activity is completely absent in *Df(btd,Sp1)* mutant embryo. The remaining *Dll* expression is likely derived from a *Dll* early enhancer. (D-G) *Df(btd,Sp1)* stage 14 mutant embryos where the expression of UAS-*Sp1* (D), UAS-*Sp1*^*ΔBTD-box*^ (E), UAS-*btd* (F), UAS-*btd* ^*ΔBTD-box*^ (G) is induced in the second thoracic segment with the *prd-Gal4* line. Note the rescue of *Dll* expression and *Dll-LT* activity (arrow) in all the genetic combinations.

**Figure S3: Leg phenotypes caused by the misexpression of *Sp1* and *btd* without the BTD box.** Adult leg phenotypes caused by the misexpression of *Sp1*, *Sp1*^*ΔBTD-box*^, *btd* and *btd* ^*ΔBTD-box*^ with the *Dll-Gal4* line. Note that the leg truncations caused by the expression of the UAS versions of *btd* and *Sp1* without the BTD box extended less distal than the wild type versions (brackets). cox, coxa; f, femur; tar, tarsus; tb, tibia; tr, trochanter.

**Figure S4: Identification of the *danr1* CRM.** Schematic of the *dan* and *danr* genomic locus in which open chromatin regions, identified by FAIRE seq for eye-antenna and leg imaginal discs are indicated by orange peaks. Data obtained from McKay and Lieb (2013) [62]. In the upper part of the panel, horizontal bars represent the DNA elements for which Gal4 drivers were generated by the VDRC (blue bars) and the Janelia Farm consortium (orange bars). None of those lines reproduced *dan* or *danr* expression in the antenna. Green bars represent DNA regions with differential open chromatin profiles between the eye-antenna and leg disc that have been selected for cloning into a reporter GFP construct. Only the danr1 fragment faithfully reproduced the expression of *dan/danr* in antenna and leg imaginal disc. Mutation of the 8 Sp1 putative binding sites slightly derepresses *danr1* activity in the leg disc (arrow).

